# Continuous Prediction of Mice Lever-Pressing Kinematic Parameters by Background Removed Single-photon Calcium Images

**DOI:** 10.1101/2024.12.17.628875

**Authors:** Mingkang Li, Ruixue Wang, Guihua Wan, Yuqi Yang, Shaomin Zhang

## Abstract

Calcium imaging has gained extensive application in neural decoding tasks because of its high precision in observing cortical neural activity. Nevertheless, the immense data volume and complexity of automated signal extraction algorithms in calcium imaging result in significant delays in extracting neuronal calcium fluorescence signals, greatly constraining the efficiency of neural decoding research and its applicability in real-time tasks. Although a few studies have successfully used partial neuronal signals from calcium imaging data for real-time neural decoding and brain-computer interface tasks, they fail to leverage the complete neuronal dataset from experiments, which limits their ability to decode continuous and complex movements. In response to this challenge, we introduce a neural decoding method based on background-removed single-photon calcium images. This approach extracts three-dimensional spatiotemporal representations of neuronal activity via background removal and employs a decoder combining 3D-ResNet and RNN networks to enable continuous and rapid decoding of mouse lever-pressing kinematic parameters. Compared with traditional methods for neural decoding using single-photon calcium imaging, this approach offers higher accuracy and faster speed. Combined with real-time motion correction algorithms, the proposed neural decoding approach meets real-time decoding requirements at a 20Hz acquisition frame rate, achieving single decoding in just 21.8ms. This advancement significantly improves the efficiency of single-photon calcium imaging-based neural decoding, offering solutions for its application in real-time tasks, such as optical brain-computer interfaces.

## 1. Introduction

Calcium imaging has been widely applied to observe neural circuits in the brain, establishing itself as a vital tool in neuroscience research^[1,2,3,4]^. Neuroscientists have utilized calcium imaging technology to obtain dynamic calcium fluorescence from neurons, allowing for the decoding of forelimb movements^[5]^, spatial locations^[6,7]^, and auditory and visual sensory information^[8,9,10]^. However, these studies depend on offline processing^[11,12,13,14,15]^ of the acquired data to extract neuronal calcium fluorescence activity, which is subsequently fed into a decoder^[16, 17]^. The data processing involved in calcium imaging typically demands considerable time. This is partly attributed to the large data volume generated by calcium imaging, which can amount to 100 GB per hour^[18]^, manifesting as images and videos. Additionally, calcium imaging data encompasses a considerable amount of background fluorescence, blood vessels, and noise; the calcium fluorescence signals from neurons are generally sparsely distributed within the field of view(FOV), complicating their extraction^[19]^. This often requires computationally intensive steps or complexity of the algorithms to improve accuracy, resulting in further time expenditure^[20]^. Such circumstances significantly restrict the efficiency of neural decoding research utilizing calcium imaging techniques and hinder the application of calcium imaging technology in real-time contexts, including brain-computer interfaces.

In the context of rapid processing of calcium imaging data, several studies have successfully implemented online motion correction, with some achieving operation in real-time scenarios. NoRMCorre^[21]^ offers an online non-rigid motion correction framework specifically designed for calcium imaging data. Mitani et al.^[22]^ have proposed a real-time motion correction solution tailored for two-photon calcium imaging. Additionally, Li et al.^[23]^ introduced a real-time motion correction plugin for single-photon calcium imaging data, which can be integrated with the Miniscope^[38]^ to facilitate real-time motion correction for each acquired frame. Collectively, these findings indicate that current research has effectively addressed the processing demands of motion correction in real-time environments.

However, with respect to automatic signal extraction in calcium imaging, although current research has introduced several rapid processing strategies^[24,25,26,27]^, the processing speed of these approaches remains relatively constrained. Currently, the predominant automatic signal extraction algorithm is based on non-negative matrix factorization^[28,29]^, which models calcium imaging data and separates spatial and temporal components indicative of neuronal activity. Algorithms grounded in non-negative matrix factorization are now extensively employed in the signal extraction for calcium imaging data. Nonetheless, these techniques generally necessitate considerable computational time and are tailored for offline processing, thereby limiting their applicability to closed-loop tasks with real-time demands. Although certain studies have improved processing speeds by implementing image scaling and reducing iterations^[24,25]^, these approaches generally cannot both meet the requirement of accuracy and computational resources^[20]^.As a result, these algorithms are still inadequate for real-time experimental processing.

In order to mitigate the time consumption during neuron signal extraction, some studies have chosen to pre-label a portion of neurons and extract their signals in real time for decoding or other experiments^[29, 30]^. While this approach substantially enhances processing speed, it discards a considerable amount of spatial information inherent in the calcium imaging data, thereby failing to leverage the full potential of calcium imaging. Other research endeavors have chosen to decode directly from calcium imaging data, utilizing two-photon calcium imaging images to perform classification tasks^[31,32]^. This approach involves projecting a sequence of two-photon calcium imaging images along the temporal axis to produce a two-dimensional image, which is subsequently used as input for the decoder, thus circumventing the signal extraction step and completing the behavior classification task. However, it is unfortunate that these studies have not addressed more complicated regression tasks, such as decoding the trajectory of forelimb movements or pressure values in lever-pressing experiments. In conclusion, current studies have not yet enabled the use of calcium imaging for continuous and complex neural decoding tasks in real-time scenarios, making it difficult to apply calcium imaging in real-time closed-loop tasks and restricting its development.

In response to this issue, we introduce an online neural decoding approach using single-photon calcium imaging data, employing these images directly as input features to predict the lever-pressing kinematic parameters. By eliminating background fluorescence and noise from the FOV, we acquire data that approximates three-dimensional spatiotemporal neuronal information. This forms the basis for a scheme that enables continuous, rapid decoding of pressure values during mouse lever-pressing, using background-removed images as samples. Subsequently, online decoding of the samples is conducted using a decoder integrating a 3D residual convolutional neural network and a recurrent neural network. This approach not only circumvents the intricate calcium signal extraction procedure, thereby conserving significant computational resources throughout the decoding process, but also utilizes fluorescent activity data from calcium imaging, minimizing the loss of signals pertinent to the behavior. Relative to traditional calcium imaging-based decoding workflows, this method demonstrates enhancements in both processing speed and accuracy, thereby fulfilling the performance demands of real-time applications. This method supports real-time closed-loop experiments and optical brain-machine interface technology using single-photon calcium imaging, broadening the applications of single-photon calcium imaging technology.

## 2. Methods

### 2.1 Motivation of decoding by calcium images

To explain the feasibility of decoding using single-photon calcium imaging images, we use the model of single-photon calcium imaging as an example. The expression for this model is as follows^[29]^:

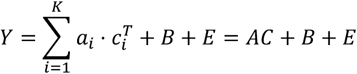

In this model, 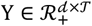 denotes the observed single-photon calcium imaging data, 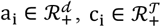 indicate the spatial and the temporal vector of the neuron *i* respectively, where *d* is the number of pixels in the FOV and *T* represents the number of the frames. *K* indicates the number of the neurons in the FOV. A = [a1, …, a_k_] and C = [c_1_, …, c_k_]^T^ represent the spatial and temporal matrix of the neurons, 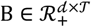 denotes background fluorescence, while 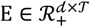 represents noise. In traditional calcium imaging-based decoding tasks, signal extraction algorithms isolate the temporal component *C* from the raw data *Y*. However, the presence of multiple unknown variables in the model makes the loss function non-convex. To address this, the problem is typically reformulated by solving for one unknown matrix at a time, treating the other variables as constants. This alternating estimation method reduces the loss function but remains computationally intensive and time-consuming, hindering neural decoding efficiency.

In decoding tasks, the temporal component *C* is generally used as the decoder’s input(Figure 1A). However, given the complexity of solving *C* through matrix decomposition, we propose bypassing this step by directly using the three-dimensional spatiotemporal signals, denoted as *AC*, as input for decoding. Theoretically, *AC* retains all the information that *C* provides, making it available approach for neural decoding. Therefore, our objective is to extract the 3D spatiotemporal signals *AC* as input features for decoding. To achieve this, we employed the background removal method^[11]^, using anisotropic diffusion filtering^[33]^ and morphological opening^[34]^ to remove background fluorescence *B* and noise *E*, producing image data that closely approximates *AC*.

**Figure 1.**
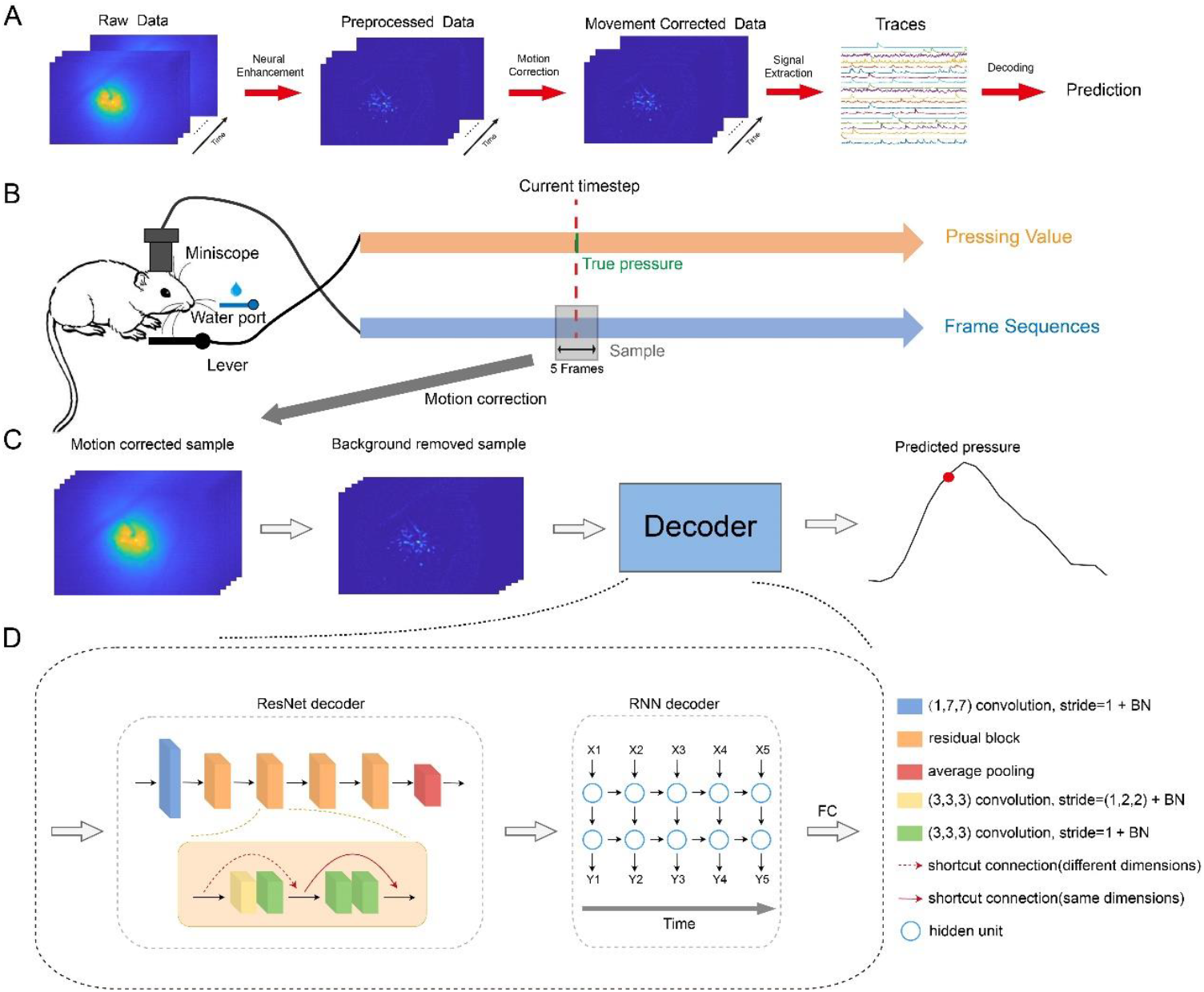
A: Example of traditional decoding workflow using single-photon calcium imaging data. The calcium imaging data processing pipeline in the example is MIN1PIPE. B: A framework for decoding lever-press behavior in mice using single-photon calcium imaging data. For each time point in the lever-pressing process, we employ the corresponding single-photon calcium imaging frame, along with the two adjacent frames before and after, to create a decoding sample. C: Decoding procedures by using single-photon calcium images. All images will undergo motion correction, followed by background removal, and then be input into the decoder to predict the pressure value at the current time point. D: Design of the decoder. The decoder consists of a three-dimensional residual network (3D-ResNet) in series with a recurrent neural network (RNN).

### 2.2 Experimental data and decoding workflow

Subsequently, we chose the lever-press behavior in mice as our research paradigm ^[37]^. Briefly, the mouse is placed in a behavior chamber allowing free movement. Upon the start of an auditory cue, the mouse begins to press the lever. If the pressure value surpasses a set threshold and sustains for a certain period, the press is deemed successful, and the mouse receives a reward; otherwise, the attempt fails, and no reward is given. In this manner, we employed Miniscope^[38]^ to synchronously collect single-photon calcium imaging data from layers 2/3 of the mouse motor cortex during the lever-press process. The surgery procedures and equipment of data acquisition were described in the previous study^[44]^. The frame rate of acquisition is 20 Hz. We collected a total of six sessions of data from three mice.

To decode pressure values during mouse lever-pressing, we used kinematic data to identify the peak pressure point in each successful lever-press event. We selected 9 frames before and 10 frames after the peak, totaling 20 frames, to represent the lever-pressing period. Additionally, we designed the decoding scheme illustrated in Figures 1B and 1C. Pressure value prediction at each time point was based on a sliding window of three-dimensional single-photon calcium images. Compared to single-frame images, the three-dimensional images within the sliding window provide additional temporal information. In addition, Given the brief delay in calcium indicator responses to action potentials, where fluorescence intensity gradually increases, we employed a sliding window of 5 frames, capturing data from the current time point and the two frames before and after. This approach more effectively captures neuronal activity during lever-pressing.

### 2.2 Design of the decoder

To process three-dimensional image data, we developed a decoder (Figure 1D) designed to predict the pressure values for each sample. The decoder consists of a three-dimensional residual network (3D-ResNet) in series with a recurrent neural network (RNN). The 3D-ResNet compresses the spatial dimensions of the input samples and extracts spatial features, producing an intermediate variable representing hidden states. This intermediate variable is then fed into the RNN to extract temporal features, and the final predicted pressure value is output through a fully connected layer.

The design of this decoder draws inspiration from conventional calcium imaging decoding workflows, which use automated matrix decomposition algorithms to identify and extract neuronal calcium fluorescence signals in the FOV, and then use these signals to decode the animal’s kinematic parameters. Typically, decoders like RNNs or other models suitable for temporal decoding are employed. Similarly, the 3D-ResNet extracts low-dimensional temporal variables from the three-dimensional image samples, which are then input into the RNN for pressure prediction. However, during the mouse lever-pressing process, not all fluorescence activity in the FOV relates to the pressing behavior. Some neurons exhibit spontaneous dynamics related to basic physiological functions, while others respond to activity from neighboring cortical neurons^[35]^. Consequently, the extracted calcium fluorescence signals often contain information unrelated to kinematics. This decoder is specifically designed to identify kinematics-related spatial features from the input three-dimensional image data. Through supervised training, the 3D-ResNet transforms kinematics-related fluorescence features into low-dimensional temporal hidden variables, enabling the RNN to predict pressure values from these variables. This approach bypasses the complex signal extraction step and directly captures task-relevant information, significantly improving decoding efficiency.

The learning rate of the model was set to 3×10−^6^, with a total of 120 epochs, and the loss function applied was mean squared error(MSE). The optimizer used for the model was the adaptive moment estimation(Adam)^[45]^.

### 2.3 Evaluation of decoding accuracy

To evaluate the performance of our proposed method, we followed the conventional calcium imaging decoding process, applying two offline signal extraction methods, MIN1PIPE^[11]^ and EXTRACT^[36]^, to process the collected calcium imaging data. These methods extracted calcium fluorescence traces from neurons in each dataset. The datasets were generated similarly, with the only difference being the application of the sliding window to the calcium fluorescence traces for sampling. Additionally, an RNN was used to decode the calcium fluorescence trace samples. This RNN had the same structure as the one described in Figure 1D, with the only difference being that the input channel count corresponded to the number of neurons in the calcium fluorescence trace samples.

For each decoding method, we employed five-fold cross-validation for accuracy assessment. Each set of samples was split into five equal parts, with three used as the training set, one as the validation set, and one as the test set. Once the model had completed the specified number of training epochs, we selected the parameters of the model with the best performance on the validation set (i.e., the lowest MSE). Finally, these parameters were used to evaluate the decoding accuracy in the test set.

### 2.4 Evaluation of decoding speed

To compare the time required by our method with traditional decoding workflows, we selected 10 single-photon calcium imaging videos, each containing 1000 frames, to assess the processing time for MIN1PIPE and EXTRACT. The extracted calcium fluorescence signals were used to generate decoding samples, and the time required for the RNN to process 1000 calcium fluorescence samples was recorded. Notably, these samples were used solely to test algorithm runtime and did not contain corresponding kinematic data. The combined time for signal extraction and RNN processing represents the total decoding time in traditional workflows. Similarly, 10 sets of image samples, each containing 1000 images, were generated from the same 10 videos, and the time required for processing these sets with the proposed decoder was recorded to represent the runtime of the image-based decoding method.

MIN1PIPE focuses on precise neuronal signal extraction in offline scenarios, incorporating complex features and algorithms, which results in relatively longer processing times. In contrast, EXTRACT uses fewer processing steps and leverages GPU acceleration to expedite computations, leading to faster processing times. Both methods rely on matrix decomposition for neuronal signal extraction, meaning that the number of neurons determines the matrix dimensions after decomposition, which directly impacts the extraction time. Therefore, we recorded the processing time for each algorithm based on datasets with varying neuron counts. All tests were conducted on a machine equipped with an Intel Core i7-8700 processor, 64 GB of RAM, and an NVIDIA GeForce RTX 3090 TI GPU.

### 2.5 Dimensionality reduction and visualization for the output of ResNet

The design of this decoder draws from traditional calcium imaging decoding approaches, but our method focuses primarily on extracting kinematics-related information from the FOV. Specifically, the 3D-ResNet replaces the signal extraction algorithms by performing both feature dimensionality reduction and extraction. To validate this, we extracted the intermediate variables generated by the 3D-ResNet within the decoder and applied dimensionality reduction. For dimensionality reduction, we applied t-distributed stochastic neighbor embedding (t-SNE)^[39]^. The t-SNE algorithm excels at mapping high-dimensional data into a low-dimensional space while preserving the original data distribution, ensuring that positional relationships in the high-dimensional space remain intact after reduction.

### 2.6 Evaluation of the decoding performance of the false positive ROIs

While background removal algorithms can eliminate most of the background fluorescence in single-photon calcium imaging, background removed images often display fluorescence that deviates notably from typical neuronal shapes within the FOV. These fluorescence signals are typically regarded as false-positive neuronal signals^[40]^. Most signal extraction algorithms incorporate a final discrimination step to filter out such false positives^[11,13,14]^, while some studies introduce manual screening to better align the extracted signals with actual neuronal dynamics^[41]^. However, such unexplained fluorescence is often due to local background fluctuations caused by out-of-focus neuronal activity, which may still be kinematically relevant. Therefore, in decoding tasks, these fluorescence signals may not always require removal.

To evaluate this hypothesis, we adjusted the neuron size parameter in MIN1PIPE, which defines the assumed diameter of neuronal cell bodies in the algorithm, and extracted calcium fluorescence matching this defined size. Reducing this parameter allowed the algorithm to more readily extract smaller fluorescence signals in the FOV. Additionally, we disabled the false-positive discrimination step in MIN1PIPE, enabling the algorithm to extract as much fluorescence as possible. Finally, we used the extracted calcium fluorescence traces, including false-positive signals, to generate samples for decoding and compared their performance to image-based decoding and decoding based on MIN1PIPE-extracted calcium trace samples.

### 2.7 Validation of the real-time decoding performance of decoding by calcium images

To evaluate the performance of our proposed decoding algorithm in real-time scenarios, we simulated data input in an online process using a real-time motion correction algorithm^[23]^ to process the data. We also implemented the background removal method in PyTorch and utilized GPU acceleration for faster processing. We then applied the decoding scheme from Section 2.1 for online neural decoding. As a comparison, we used offline-extracted neuronal spatial contours to extract calcium fluorescence traces during the online process and performed online decoding. Additionally, we employed the real-time decoding method based on manually labeled neurons proposed by Liu et al. as another control group. Throughout this process, we recorded the time consumed at each step and evaluated the accuracy of the different methods in a simulated online setting.

## 3 Results

### 3.1 The accuracy of decoding by calcium images is superior to that of decoding by traces

After successfully training three mice for lever-pressing tasks, we collected single-photon calcium imaging and corresponding pressure value data from six lever-press tasks. The number of neurons identified using MIN1PIPE and EXTRACT through offline analysis varied, as did their spatial distribution (Figure 2A). Decoding using different sample types revealed that pressure values decoded from image data were closer to actual values (Figure 2B). In all six datasets, the correlation coefficients (CC) for image samples decoding was significantly higher than that for calcium fluorescence trace data decoding (Figure 2C) (Image samples vs. trace samples (MIN1PIPE), p < 0.05, Wilcoxon signed-rank test, n = 6; Image samples vs. trace samples (EXTRACT), p < 0.05, Wilcoxon signed-rank test, n = 6). Additionally, the mean squared error (MSE) from image-based decoding was significantly lower (Figure 2D) (Image samples vs. trace samples (MIN1PIPE), p < 0.05; Image samples vs. trace samples (EXTRACT), p < 0.05; Wilcoxon signed-rank test, n = 6).

**Figure 2.**
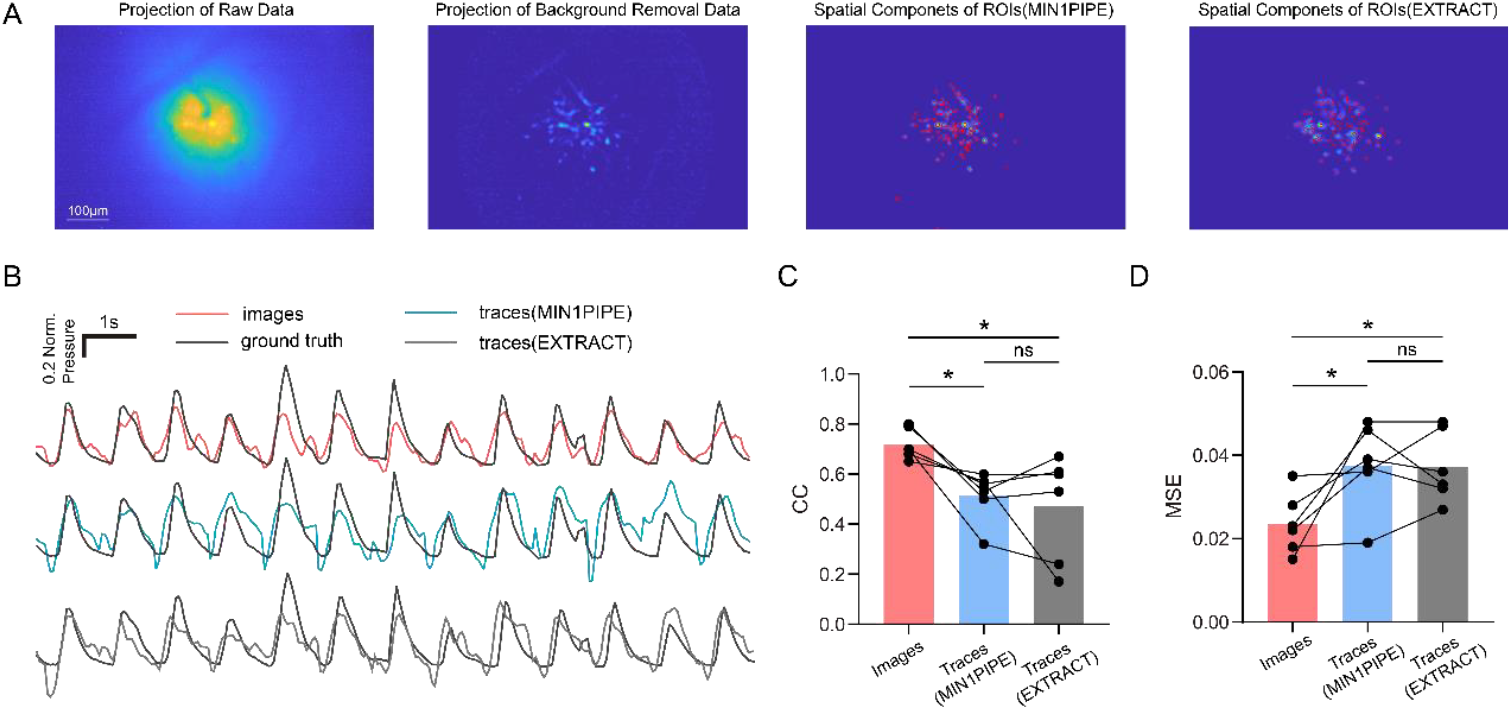
A: Projection image of raw data and background removal data and spatial footprints of the ROIs extracted by MIN1PIPE and EXTRACT. B: Example of the predicted pressing values (red: decoding by images; cyan: decoding by traces extracted by MIN1PIPE; gray: decoding by traces extracted by EXTRACT) and the ground truth(black). C: Correlation coefficients between the predicted values and the true values from 6 sessions(images vs. traces(MIN1PIPE): *p<0.05, Wilcoxon signed-rank test, n=6, images vs. traces(EXTRACT): *p<0.05, Wilcoxon signed-rank test, n=6, traces(MIN1PIPE) vs. traces(EXTRACT): p>0.05, Wilcoxon signed-rank test, n=6). D: Mean squared errors between the predicted values and the true values from 6 sessions(images vs. traces(MIN1PIPE): *p<0.05, Wilcoxon signed-rank test, n=6, images vs. traces(EXTRACT): *p<0.05, Wilcoxon signed-rank test, n=6, traces(MIN1PIPE) vs. traces(EXTRACT): p>0.05, Wilcoxon signed-rank test, n=6).

The number and spatial location of neurons identified by the two offline signal extraction algorithms in each dataset indicate that different algorithms result in varying neuron extraction outcomes. These differences impact the accuracy of neuron extraction in traditional decoding workflows, leading to variations in decoding performance. Although no significant difference in decoding accuracy was observed between samples extracted by MIN1PIPE and EXTRACT (p > 0.05, Wilcoxon signed-rank test, n = 6), variations in CC and MSE were noted across decoding tasks. However, when decoding by the image samples, the variability in neuron signal extraction caused by different algorithms is substantially minimized, thus preventing instability in decoding performance. By bypassing the complex signal extraction step, our method significantly reduces the influence of data processing on neuron signal extraction. Thus, the proposed decoding strategy and decoder design in our study demonstrate higher accuracy compared to traditional calcium imaging decoding workflows. Furthermore, it eliminates discrepancies in neuron extraction between different signal extraction algorithms, demonstrating that single-photon calcium imaging data can be directly applied in neural decoding tasks, especially in regression tasks like kinematic trajectory fitting—a capability not previously reported in the literature.

### 3.2 The speed of decoding by calcium images is faster and more stable

Figure 3A shows the time required to decode 1000 samples with image-based decoding, compared to the time taken by MIN1PIPE and EXTRACT to process 1000 video frames and decode 1000 calcium trace samples. Ten video datasets, each containing different numbers of neurons, were used for testing. Decoding 1000 samples from image samples took approximately 7 seconds, whereas signal extraction and decoding with EXTRACT required between 58 and 233 seconds for the same number of samples. MIN1PIPE took significantly longer, averaging 903 seconds to complete the same task. In contrast, the image-based decoding method demonstrated a significant speed advantage (ResNet+RNN vs. EXTRACT+RNN, p < 0.01, Wilcoxon signed-rank test, n = 10; ResNet+RNN vs. MIN1PIPE+RNN, p < 0.01, Wilcoxon signed-rank test, n = 10). Although EXTRACT utilizes GPU acceleration to expedite matrix decomposition, variability in the number of neurons in the FOV and differences in fluorescence signal distribution across datasets still lead to considerable fluctuations in processing speed.

**Figure 3.**
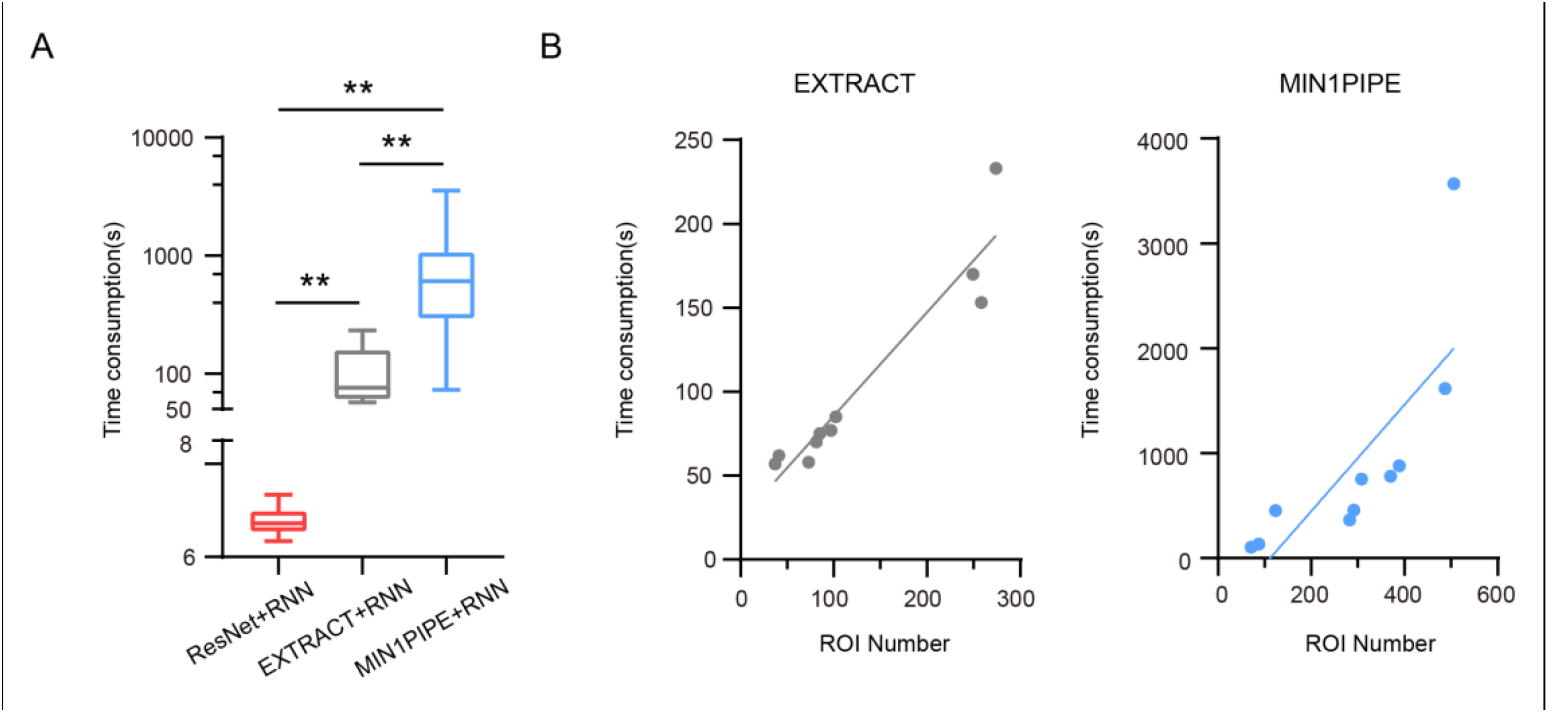
A:Time consumption of decoding 1000 images samples by ResNet+RNN decoder and time consumption of processing 1000 frames calcium imaging data by signal extraction algorithms and decoding 1000 traces samples(images vs. traces(MIN1PIPE): **p<0.01, Wilcoxon signed-rank test, n=10; images vs. traces(EXTRACT): **p<0.01, Wilcoxon signed-rank test, n=10; traces(MIN1PIPE) vs. traces(EXTRACT): **p<0.05, Wilcoxon signed-rank test, n=10). B: The variation in time consumption of the signal extraction algorithms (MIN1PIPE and EXTRACT) when processing 1000 frames data as a function of the number of extracted ROIs. The lines indicate linear fitting.

To further assess the impact of neuron count on the processing speed of signal extraction algorithms, we conducted linear fitting of the processing time for MIN1PIPE and EXTRACT against the number of neurons identified in each dataset (Figure 3B). The fitting results showed that the processing time for both methods was affected by the number of neurons in the FOV. As the number of detected neurons increased, the processing time increased accordingly, though to varying degrees. For EXTRACT, the processing time ranged from 58 seconds to a maximum of 233 seconds, while for MIN1PIPE, it increased from 73 seconds to up to 3570 seconds. This phenomenon occurs because these automated signal extraction algorithms decompose raw data into two matrices representing temporal and spatial components, with one matrix’s dimensions determined by the number of detected neurons. As the number of detected neurons increases, the dimensions of these matrices expand, increasing the computational load required to minimize the loss function. In contrast, image-based decoding does not face this issue, as the processing time for decoding three-dimensional image data depends only on data size, regardless of the number of neurons in the FOV. This advantage enables image-based decoding to maintain relatively stable processing speeds across samples from different time points, brain regions, and experimental animals, which is crucial for applying calcium imaging technology in real-time settings like brain-machine interfaces.

### 3.3 Image decoding enables better extraction of kinematics-related features within the FOV

Figure 4 presents the t-SNE projection of the intermediate hidden variables output by the 3D-ResNet and the calcium trace samples onto a two-dimensional plane, with each sample labeled based on its corresponding pressure value and temporal index. A subset of data from three mice was selected for visualization, with each sample marked by its actual pressure value and temporal index.

**Figure 4.**
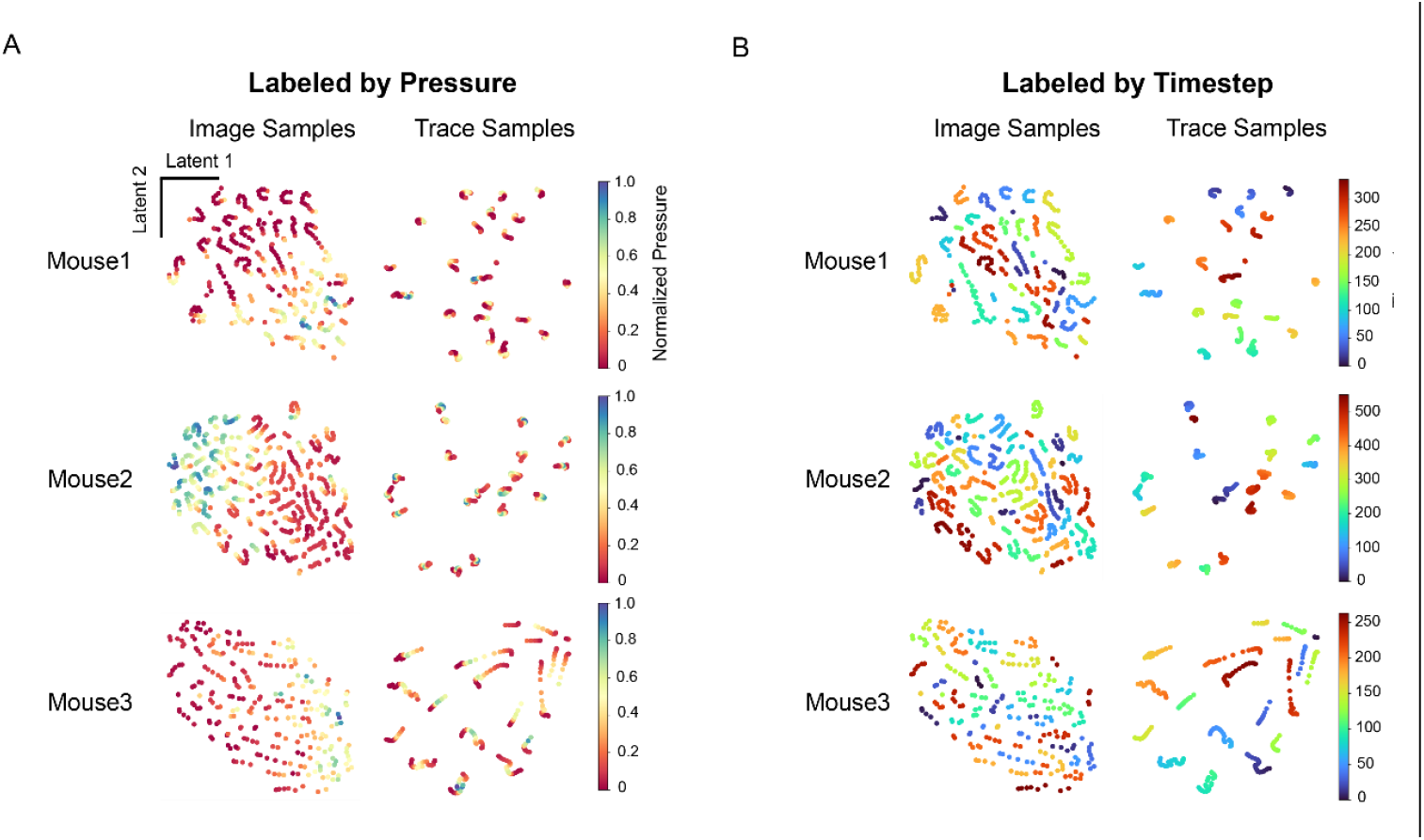
Dimensionality reduction of the outputs from ResNet decoder when decoding by calcium images and the calcium trace samples is performed using the t-SNE algorithm, with each row representing a subset of samples from a single mouse. The outputs of ResNet show a correlation with pressure values in low-dimensional space. A: Using normalized pressure as labels, the dimensionally reduced samples are annotated. The distribution of intermediate variables from image decoding in low-dimensional space correlates with pressure values, with a diagonal pattern of high and low pressure. Calcium trace samples, however, do not display this pattern in low-dimensional space. B: The dimensionally reduced samples are labeled by time steps. Temporally adjacent calcium trace samples are clustered in low-dimensional space, with each cluster representing samples from a single lever-press period. However, for the outputs from ResNet, even temporally close samples are separated because of the differences in the pressure values they represent.

After dimensionality reduction, the intermediate variables from the 3D-ResNet output were divided into two distinct regions based on pressure values (Figure 4A). Samples with high pressure values predominantly clustered on one side along the diagonal of the two-dimensional space, while those with low pressure values were distributed on the opposite side. In contrast, calcium trace samples did not exhibit any clear distribution pattern related to pressure values after dimensionality reduction. These samples formed multiple clusters, with each cluster containing both high– and low-pressure value samples. When the reduced-dimension samples were re-labeled by temporal index(Figure 4B), calcium fluorescence trace samples clustered by their temporal sequence, indicating a strong temporal correlation. This suggests that the signal extraction algorithm primarily captured temporal information and inter-neuronal correlations in the original data, but showed limited association with kinematics.

Additionally, after mapping to the low-dimensional space, some calcium fluorescence trajectory samples formed tight clusters, dividing into distinct categories. This clustering occurred because we selected only calcium imaging and kinematic data from the lever-pressing tasks, with each category representing samples from the corresponding lever-pressing period. These samples clustered together in the low-dimensional space due to their temporal proximity. In contrast, the temporal correlation among the intermediate variables output by the 3D-ResNet was much weaker. Temporally adjacent samples were mapped to distant locations because of differences in their corresponding pressure values. This observation supports our hypothesis in decoder design, indicating that the 3D-ResNet effectively extracts kinematics-related features from the data and generates corresponding temporal hidden variables, improving the efficiency of feature extraction in the subsequent RNN network. These results further demonstrate the capability of single-photon calcium imaging data to be directly applied in neural decoding.

### 3.4 The background components of single-photon calcium imaging images also contribute to decoding

After adjusting the parameters of MIN1PIPE and eliminating the false-positive neuron screening step, the spatial distribution of extracted neurons changed (Figure 5A), and the number of detected neurons increased (Figure 5B). The algorithm additionally detected numerous low-intensity fluorescence signals scattered around the periphery of the FOV, which clearly lacked the contours of real neurons. In contrast, data processed with MIN1PIPE’s recommended parameters and false-positive screening steps more accurately reflected the spatial characteristics of real neurons and exhibited stronger fluorescence signals.

**Figure 5.**
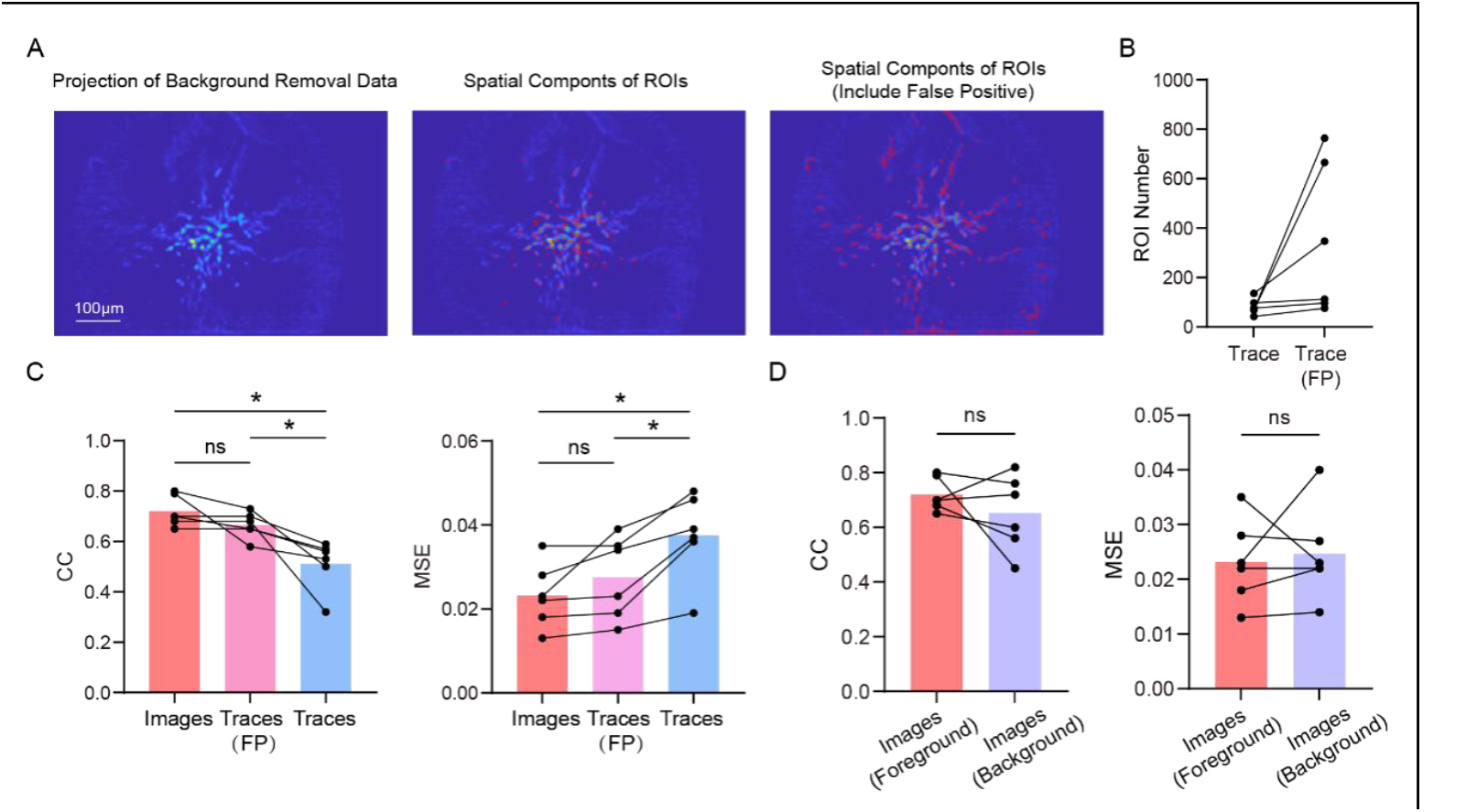
A: Projection image of the background removal data(left) and spatial footprints of the ROIs(middle) and the ROIs including false positive (FP) signals(right) extracted by MIN1PIPE. B: Extracted ROI number (ROIs vs. ROIs including FP signals). C: Decoding accuracy of image samples, trace samples including FP signals and trace samples(CC and MSE: images vs. traces(FP): p>0.05, Wilcoxon signed-rank test, n=6; images vs. traces: *p<0.05, Wilcoxon signed-rank test, n=6; traces(FP) vs. traces: *p<0.05, Wilcoxon signed-rank test, n=6). D: Decoding accuracy of foreground images and background images(CC and MSE: images(foreground) vs. images(background): p>0.05, Wilcoxon signed-rank test, n=6).

Figure 5C shows the decoding performance of samples that include false-positive neurons. No significant differences were found between the performance of false-positive neuron samples and image samples (CC: p > 0.05, Wilcoxon signed-rank test, n = 6; MSE: p > 0.05, Wilcoxon signed-rank test, n = 6), though the mean CC was higher and mean MSE was lower for image samples.

Additionally, the decoding performance of samples with false-positive neurons showed a significant improvement in CC compared to the pre-adjustment performance (p < 0.05, Wilcoxon signed-rank test, n = 6), along with a significant reduction in MSE (p < 0.05, Wilcoxon signed-rank test, n = 6). These results suggest that, although the added false-positive neurons lacked the prominent characteristics of real neurons, these abnormal signals, likely caused by background fluorescence fluctuations, are kinematically relevant and should be considered in decoding tasks. When using single-photon calcium images for decoding, the sample data contained both the three-dimensional spatiotemporal information of real neurons and the dynamic activity of these false-positive signals, resulting in better decoding performance.

Additionally, we posited that background fluorescence may carry kinematics-related information. To test this, we used background components extracted during the removal process to generate corresponding decoding samples and performed the same neural decoding validation. The decoding accuracy using background images is shown in Figure 5D. All background samples exhibited decoding ability, indicating that background fluorescence in single-photon calcium imaging data also carries kinematics -related information. However, compared to foreground image decoding, background-based decoding showed more significant fluctuations in performance. In some datasets, background decoding outperformed foreground decoding, but in others, it performed worse. This variability occurs because experimenters cannot easily determine which depth contains the most kinematically relevant neurons during imaging, and single-photon imaging captures fluorescence fluctuations from out-of-focus neurons in the background. When out-of-focus neurons are more kinematically relevant, background fluorescence fluctuations may lead to higher decoding accuracy; conversely, when focal plane neurons contain more kinematically relevant information, foreground decoding performs better. Despite the variability in decoding accuracy across datasets, foreground decoding exhibited greater stability. Thus, foreground images, representing the three-dimensional spatiotemporal information of neurons, should be prioritized as inputs for decoding.

### 3.5 Decoding using images can meet the demands of real-time applications

Figure 6A presents the accuracy results of the proposed decoding scheme in a simulated online scenario. The online decoding method based on single-photon calcium images achieved the highest accuracy (CC and MSE: image samples vs. trace samples, p < 0.05, n = 6, Wilcoxon signed-rank test; image samples vs. manually labeled samples, p < 0.05, n = 6, Wilcoxon signed-rank test; trace samples vs. manually labeled samples, p < 0.05, n = 6, Wilcoxon signed-rank test). In contrast, the commonly used real-time decoding method, which relies on manually labeling a subset of neurons, performed poorly in the continuous pressure decoding task. This indicates that relying solely on partial neuronal information is insufficient for handling complex continuous decoding tasks and brain-machine interface (BMI) research, and is only suitable for simpler classification scenarios. As data dimensionality increases, decoding performance in the simulated online scenario significantly improves, regardless of whether offline-extracted calcium fluorescence signals or single-photon calcium imaging images were used. This further demonstrates that fully leveraging the information in calcium imaging images is essential for ensuring the accuracy of continuous motion decoding in neural tasks. Sacrificing input dimensionality for performance improvements should be avoided.

**Figure 6.**
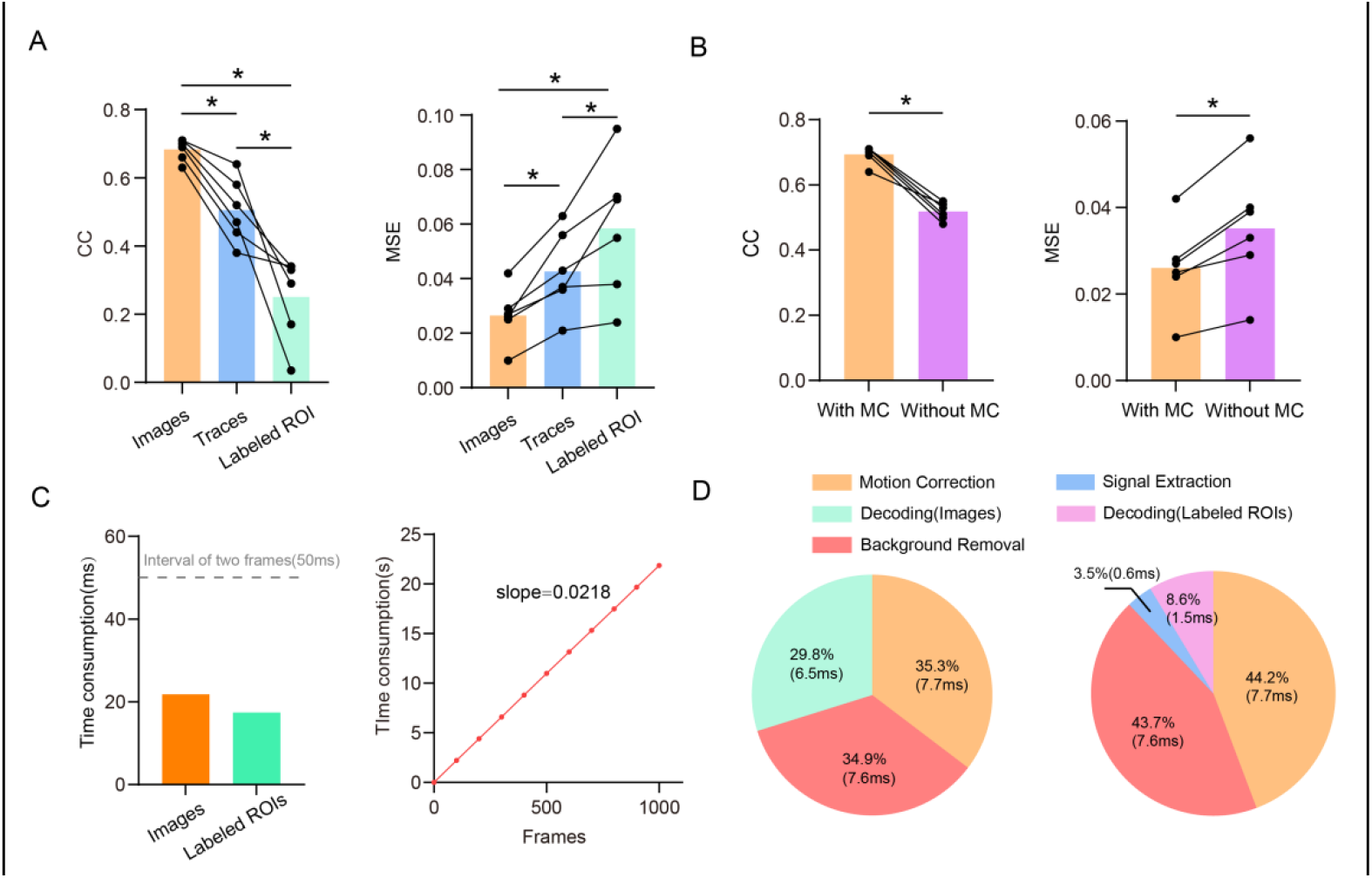
A: Decoding accuracy of online decoding by calcium images, traces extracted by MIN1PIPE and traces from 10 labeled ROIs(CC and MSE: images vs. traces, p<0.05, Wilcoxon signed-rank test, n=6; images vs. labeled ROIs, p<0.05, Wilcoxon signed-rank test, n=6; traces vs. labeled ROIs, p<0.05, Wilcoxon signed-rank test, n=6.). B: The influence of motion correction(MC) for decoding accuracy(CC and MSE: with MC vs. without MC, p<0.05, Wilcoxon signed-rank test, n=6.). C: Left: Speed of online decoding by calcium images and by 10 labeled ROIs. Time consumption of decoding a calcium image sample is 21.8ms and that of decoding a trace sample of labeled ROIs is 17.4ms. The gray dot line indicates the interval of two frames at 20Hz acquisition rate. Right: The change in decoding time using image samples as frame count increases. The cumulative time is recorded every 500 frames. The red line represents a linear fit. D: Time consumption of each procedure in decoding by calcium image sample(left) and 10 labeled ROIs(right).

Figure 6B illustrates the impact of motion correction on decoding performance. When images were subjected to random offsets of 5 to 10 pixels, decoding performance significantly declined compared to data with motion correction (p < 0.05, Wilcoxon signed-rank test, n = 6). This result indicates that when motion artifacts are present in the FOV, neuronal temporal information is disrupted, and the decoder cannot achieve high accuracy relying solely on the spatial information within the FOV. Therefore, motion correction is essential for neural decoding based on single-photon calcium imaging. While continuous movement in the FOV may not occur in real-world applications, and the extent of motion is difficult to predict, it cannot be guaranteed that experimental data used for decoding will be free from motion artifacts. To ensure decoding accuracy in real-world scenarios, it is necessary to maintain motion correction throughout the entire decoding process. Additionally, some motion in the FOV may be caused by the animal’s behavior, making it difficult for researchers to determine whether the decoder is predicting based on neuronal activity or systematic motion artifacts. Thus, even if the motion artifacts in the FOV does not negatively impact decoding, this interference should still be removed to ensure that the decoder relies solely on neuronal activity to predict kinematics parameters.

Figure 6C presents the performance of the proposed online decoding scheme. The total time required for predicting the current pressure value is only 21.8 ms, shorter than the 50 ms interval between frames at a 20 Hz sampling rate. This indicates that the system fully meets real-time processing requirements under 20 Hz acquisition conditions. Although the manual neuron labeling method is faster, with a processing time of 17.4 ms, its accuracy is insufficient for this task. Additionally, the decoding time of the proposed scheme increases linearly with the number of input frames, demonstrating that the workflow can stably handle long data streams, meeting the design requirements for online processing.

Figure 6D illustrates the time distribution of each method during image-based decoding and manually labeled neuron decoding in the simulated online scenario. As shown, manual extraction and decoding of calcium fluorescence are faster, while image-based decoding is slower, primarily due to differences in data dimensionality, as image samples contain much more data than manually labeled neuron fluorescence signals. Despite this, the difference between the two methods is less than 5 ms, and image-based decoding achieves a significant improvement in accuracy. Therefore, to meet real-time processing requirements, increasing input dimensionality and fully utilizing the high spatial resolution of calcium imaging can further enhance decoding accuracy.

## 4. Discussion

The conventional approach to neural decoding using calcium imaging data involves the extraction of neuronal calcium signals from motion-corrected image data through automated signal extraction algorithms, a method that is resource-intensive and adversely affects the utility of calcium imaging technology in real-time decoding contexts. In response to this challenge, we propose a method for neural decoding that utilizes single-photon calcium images, employing anisotropic diffusion filtering and morphological opening operations to achieve denoising and background removal, thereby obtaining images that represent the three-dimensional spatiotemporal information of neurons for decoding. By omitting the signal extraction step, this approach conserves substantial processing time. Concurrently, we developed an online decoding task designed to predict the pressing values produced by mice, as well as a decoder specifically designed for this purpose. The decoder is constructed from a series of a three-dimensional convolutional residual network and a recurrent neural network. Relative to traditional decoding procedures that utilize single-photon calcium imaging data, this method streamlines the acquisition of kinematics-related information from images via the residual network, omitting the signal extraction step and thereby enhancing computational efficiency.

Subsequently, we employed the conventional calcium imaging decoding procedure as a control group to evaluate decoding accuracy and decoding speed. Our approach exhibits enhanced accuracy and speed relative to conventional decoding workflows. In conventional decoding approaches, automated signal extraction algorithms are predicated on matrix decomposition concepts, and their computational speed is partly contingent upon the number of neurons present in the data. Consequently, this may lead to performance discrepancies when analyzing data from various brain regions, thus complicating the accurate evaluation of the algorithm effectiveness in real-time contexts. Conversely, the time taken for decoding using calcium image data is exclusively related to the size of the images, and when employing identical acquisition devices for experiments across different animals and brain regions, the decoding performance remains consistent, thus offering more dependable reference values for the performance metrics. Moreover, a dimensionality reduction analysis of the outputs from the residual network indicates that our decoder is capable of directly acquiring kinematics-related information from the calcium images for predicting pressure values, in contrast to traces derived from signal extraction algorithms, which do not adequately reflect kinematics-relevant characteristics. This finding likewise illustrates the appropriateness of employing single-photon calcium imaging data for neural decoding tasks, thereby mitigating the impact of signal extraction algorithms on decoding efficiency.

Analyzing the false positive fluorescence in the foreground alongside background fluorescence reveals that the background components of single-photon calcium imaging data harbor kinematics-related information as well; however, these components are typically disregarded or ignored in conventional decoding tasks utilizing calcium imaging. Scattering effects during single-photon imaging can introduce fluorescence from beyond the focal plane into the collected signals^[42,43]^, appearing in the data as localized background fluctuations^[29]^. Such localized background fluorescence is generated by the calcium signals of out-of-focus neurons; when these neurons demonstrate dynamic activities associated with kinematics, leveraging background images for decoding can yield high accuracy. Nevertheless, estimating and quantifying out-of-focus background fluorescence is typically problematic, leading most studies to classify it as irrelevant information and refrain from employing it^[40]^. Consequently, in the context of neural decoding tasks, we are unable to ascertain whether the background fluorescence present in the single-photon calcium imaging data holds more information relevant to kinematics than the signals from the foreground neurons. To tackle this challenge, one approach is to modify the focal plane during imaging and select neural signals at different depths for decoding trials, thereby determining the most suitable imaging depth for effective decoding. Moreover, future studies may introduce more background information to facilitate comprehensive utilization of calcium imaging data.

Finally, we combined the method introduced in this study with a real-time motion correction algorithm to develop an online decoding system, subsequently evaluating its performance through a simulated online workflow. The system fulfills the demands of real-time neural decoding tasks with respect to both accuracy and speed, requiring only 21.8 ms to predict each sample, which is less than the 50 ms interval for acquiring two frames at a 20 Hz frame rate. Our approach demonstrates the ability to perform continuous motion decoding in real-time contexts, offering technical support for optical brain-machine interface systems that utilize single-photon calcium imaging. This method offers a solution for executing more intricate real-time tasks, including sophisticated optical brain-machine interfaces.

